# Multiple Low Dose (MLD) therapy: an effective strategy to treat EGFR inhibitor-resistant NSCLC tumours

**DOI:** 10.1101/821975

**Authors:** João M. Fernandes Neto, Ernest Nadal, Salo N Ooft, Evert Bosdriesz, Lourdes Farre, Chelsea McLean, Sjoerd Klarenbeek, Anouk Jurgens, Hannes Hagen, Liqin Wang, Enriqueta Felip, Alex Martinez-Marti, Emile Voest, Lodewyk F.A. Wessels, Olaf van Tellingen, Alberto Villanueva, René Bernards

## Abstract

Targeted cancer drugs often elicit powerful initial responses, but generally fail to deliver long-term benefit due to the emergence of resistant cells^1,2^. This is thought to be the consequence of strong selective pressure exerted on the cancer drug target by a Maximum Tolerated Dose (MTD) of a drug^3–5^. We hypothesized that partial inhibition of multiple components in the same oncogenic signalling pathway might add up to complete pathway inhibition, while at the same time decreasing the selective pressure on each individual component to acquire a resistance mutation. We report here testing of this Multiple Low Dose (MLD) model of drug administration in Epidermal Growth Factor Receptor (EGFR) mutant non-small cell lung cancer (NSCLC). We show that as little as 20% of the individual drug doses required for full inhibition of cell viability is sufficient to completely block MAPK signalling and proliferation when used in 3D (RAF+MEK+ERK inhibitors) or 4D (EGFR+RAF+MEK+ERK inhibitors) combinations. Importantly, *EGFR* mutant NSCLC cells treated with EGFR inhibitors at a high dose rapidly developed resistance in vitro, but the cells treated with 3D or 4D MLD therapy did not. Moreover, NSCLC cells that had gained resistance to high dose anti-EGFR therapies were still sensitive to MLD therapy. Using several animal models, including patient derived xenografts of NSCLC tumours that are resistant to EGFR inhibitors erlotinib and osimertinib^6,7^, we found durable responses to MLD therapy without associated toxicity. These data support the notion that partial inhibition of multiple components of cancer-activated signalling pathways is difficult to circumvent and suggest that MLD therapy could deliver clinical benefit. We propose that MLD strategy could be an effective treatment option for EGFR mutant NSCLC patients, especially those having acquired resistance to even third generation EGFR inhibitor therapy.

---

Inhibition of signalling pathways that are activated by oncogenic mutations elicit therapeutic responses due to “addiction” of the cancer to the activated pathway^8^. However, in advanced cancers, development of resistance is practically inevitable due to secondary mutations that restore signalling through the drug-inhibited pathway. Such acquired resistance mutations affect either the drug target itself or components that act upstream, downstream or parallel to the activated signalling component^1, 9^. In *BRAF* mutant melanoma and NSCLC, inhibition of two components of the same oncogenic pathway (BRAF+MEK, referred to as “vertical targeting”) has been shown to provide more lasting clinical benefit compared to inhibition of only BRAF^10, 11^. More recently, both clinical^12, 13^ and pre-clinical^14^ studies have shown that inhibition of three components of the same oncogenic pathway further increases therapeutic benefit. In these scenarios the drugs are used at maximum tolerated dose (MTD). The increase in the number of drugs being used in combination is usually accompanied by an increase in toxicity and to this date virtually no studies have been done to assess the efficacy of using drugs below-MTD. In a meta-analysis of 24 phase 1 clinical studies, patients dosed below single agent MTD did almost as well those that received MTD, suggesting that most patients are over-treated with targeted agents^15^. In a preclinical model, multiple drugs used at low dose also demonstrated promising activity in ovarian clear cell carcinoma^16^. In this study we explore the use of a Multiple Low Dose (MLD) strategy in EGFR mutant NSCLC. In this approach, multiple drugs that act in the same oncogenic signalling pathway are combined at low concentration. We hypothesized that this might add up to complete pathway inhibition without causing prohibitive toxicity. Further, by using low drug concentrations, the pressure exerted on each node of the pathway should greatly diminish, reducing the selective pressure on each node and therefore diminishing the chances of acquiring resistance.

The mechanisms of resistance to EGFR inhibition (standard-of-care) in *EGFR* mutant NSCLC are well understood. We therefore compared the efficacy of MLD therapy to standard-of-care MTD therapies in this indication. We used PC9 NSCLC cells, which harbour an activating mutation in the gene encoding EGFR^17^. We used four drugs, each inhibiting a different node in the MAPK pathway: gefitinib (EGFR inhibitor), LY3009120 (pan-RAF inhibitor^18^), trametinib (MEK inhibitor) and SCH772984 (ERK inhibitor^19^), as shown schematically in Fig. 1a. We established dose-response curves for each of the four drugs using 5-day culture assays (Fig. 1b). From these data, we inferred for all 4 inhibitors the IC_20_ dose, *i.e.* a drug concentration that inhibits cell viability by 20% - henceforth referred as Low Dose (LD). To assess the efficacy of the MLD strategy we then tested the impact of all possible drug combinations of the 4 drugs at LD on cell viability (assessed by CellTiter-Blue® assay), on cell proliferation (assessed by long-term colony formation assay) and on pathway activity (measured by p-RSK levels^20^ using Western Blotting) (Figs. 1c-e). We found that PC9 cells treated with the single drugs at low dose were only minimally affected, as expected. However, some of the drug combinations showed a striking combination effect, much higher than expected based on drug additivity. In particular, the combination of RAF+MEK+ERK inhibitors at low dose (henceforth called 3D combination) and the combination of EGFR+RAF+MEK+ERK inhibitors at low dose (henceforth called 4D combination) showed an almost complete inhibition of cell viability and proliferation, along with a complete blockade of MAPK pathway signalling. Due to these notable findings we pursued the MLD study focusing mostly on the 3D and 4D combinations. To address if we could further reduce the drug concentrations, we diluted the 4D combination. When the drugs were reduced to half of the IC_20_ concentrations, the 4D combination was no longer able to achieve complete inhibition of proliferation and was similarly unable to mediate complete MAPK pathway inhibition, indicating that there is a threshold that limits efficacy (Supplemental Figs. 1a, b). Based on this, we continued our MLD studies using the IC_20_ concentrations as “Low Dose”. To make sure our findings were not drug-specific, we tested the MLD approach using different inhibitors for each of the nodes in the MAPK pathway (erlotinib as EGFRi, BGB-283 as RAFi, selumetinib as MEKi and LY-3214996 as ERKi). Supplemental Figs. 1c, d show that we obtained essentially the same synergy with these drugs in 3D and 4D combinations. This, together with the notion that each drug is used at low dose, makes it very unlikely that off target effects of the four drugs are responsible for the observed effects.

**Figure 1:**
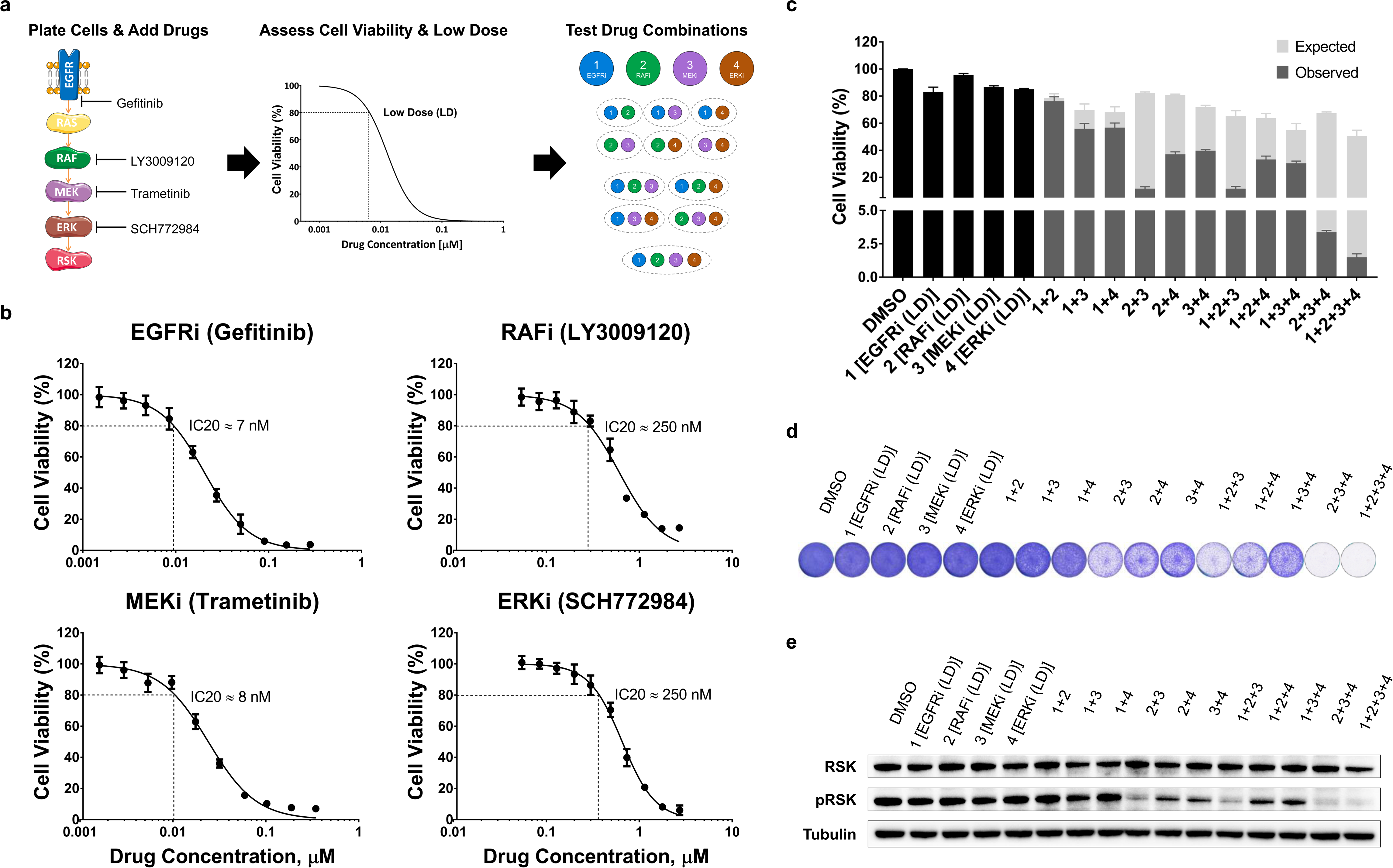
Multiple Low Dose therapy blocks MAPK pathway and proliferation in PC9 cells. **a,** xSchematic of the Multiple Low Dose (MLD) efficacy determination. After plating, cells are treated with increasing drug concentrations. Four days later cell viability is measured and the low dose (LD) is assessed. At last, the efficacy of all the possible combinations at LD is determined. **b,** Dose-response curves of EGFR, RAF, MEK and ERK inhibitors in PC9 cells. PC9 cells were cultured with increasing concentrations of EGFRi Gefitinib, RAFi LY3009120, MEKi Trametinib or ERKi SCH772984 for 4 days, after which cell viability was measured using CellTiter-Blue^®^. Standard deviation (SD) from 3 biologically independent replicates (each with 3 technical replicates) is plotted. Low doses (IC20s) were then determined: gefitinib=7nM, LY3009120=250 nM, 292 trametinib=8nM and SCH772984=250nM. **c-e,** Determination of the efficacy of all the possible combinations of EGFR, RAF, MEK and ERK inhibitors at LD in PC9 cells. PC9 cells were cultured with all possible drug combinations of EGFR, RAF, MEK and ERK inhibitors at the low doses determined in (b). In (c) cell viability was measured by CellTiter-Blue^®^ assay after 4 days of treatment; SD from 3 biologically independent replicates (each with 3 technical replicates) is plotted: in grey the observed experimental cell viability and in light-grey the expected cell viability if an additive effect was observed. In (d) cells were treated for 10 days, after which plates were stained and scanned; A representative image from the 3 biologically independent replicates performed is displayed. In (e) protein for western blotting was harvested after 24 hours of treatment; The level of pathway inhibition was determined by examining pRSK protein levels in the western blot. Tubulin was used as loading control. A representative image from the 3 biologically independent replicates performed is displayed.

Next, we tested how MLD therapy compares to standard-of-care MTD therapy in terms of resistance development. To mimic MTD therapy we treated PC9 cells with a concentration of EGFR inhibitor gefitinib that inhibited cell viability by ∼99% in a 5-day culture assay – henceforth referred as High Dose (HD). We found that 3D and 4D combinations inhibit cell proliferation and induce apoptosis at comparable levels to cells treated with HD of gefitinib (Fig. 2a and Supplemental Fig. 2a, b). The level of pathway inhibition is also similar between cells treated with 3D and 4D combinations and HD of gefitinib (Fig. 2d). Additionally, we performed RNA-Seq transcriptome analyses in cells treated with 4D combination (Supplemental Fig. 2c, d). These data showed that 4D combo treated cells displayed a significant downregulation of MYC and E2F target genes as well as cell cycle genes. Moreover, MAPK activity markers^21^ were significantly downregulated and several pro-apoptotic genes were found to be upregulated, while anti-apoptotic genes were downregulated. To study how MLD therapy compares to MTD therapy regarding resistance, we treated PC9 cells with 3D or 4D combinations and with HD of gefitinib or osimertinib for one month (Fig. 2b). As seen by others previously^22, 23^, cells treated with HD of gefitinib or osimertinib quickly developed resistance, but the cells treated with 3D or 4D combinations did not. Additionally, we treated PC9 cells for 16 days with high dose of gefitinib or with 3D or with 4D MLD combinations; we then either removed the drugs, continued to treat with the original drug, or treated with 4D MLD combination for another 16 days (Supplemental Fig. 2e). We observed resistant colonies after 32 days of gefitinib treatment, but not in the cells treated with 3D or 4D combinations. Apparently 16 day-treatment with 3D or 4D combinations had killed all cells, as continued culturing for another 16 days in media without drugs did not yield any colonies. Importantly, PC9 cells that had developed resistance to high dose EGFR inhibitor, were still responsive to 4D MLD combination. This striking result indicates that EGFR inhibitor-resistant cells remain sensitive to 3D and 4D combinations. This suggests that MLD therapy might be an option for patients having developed resistance to standard-of-care EGFR inhibitor therapy.

**Figure 2:**
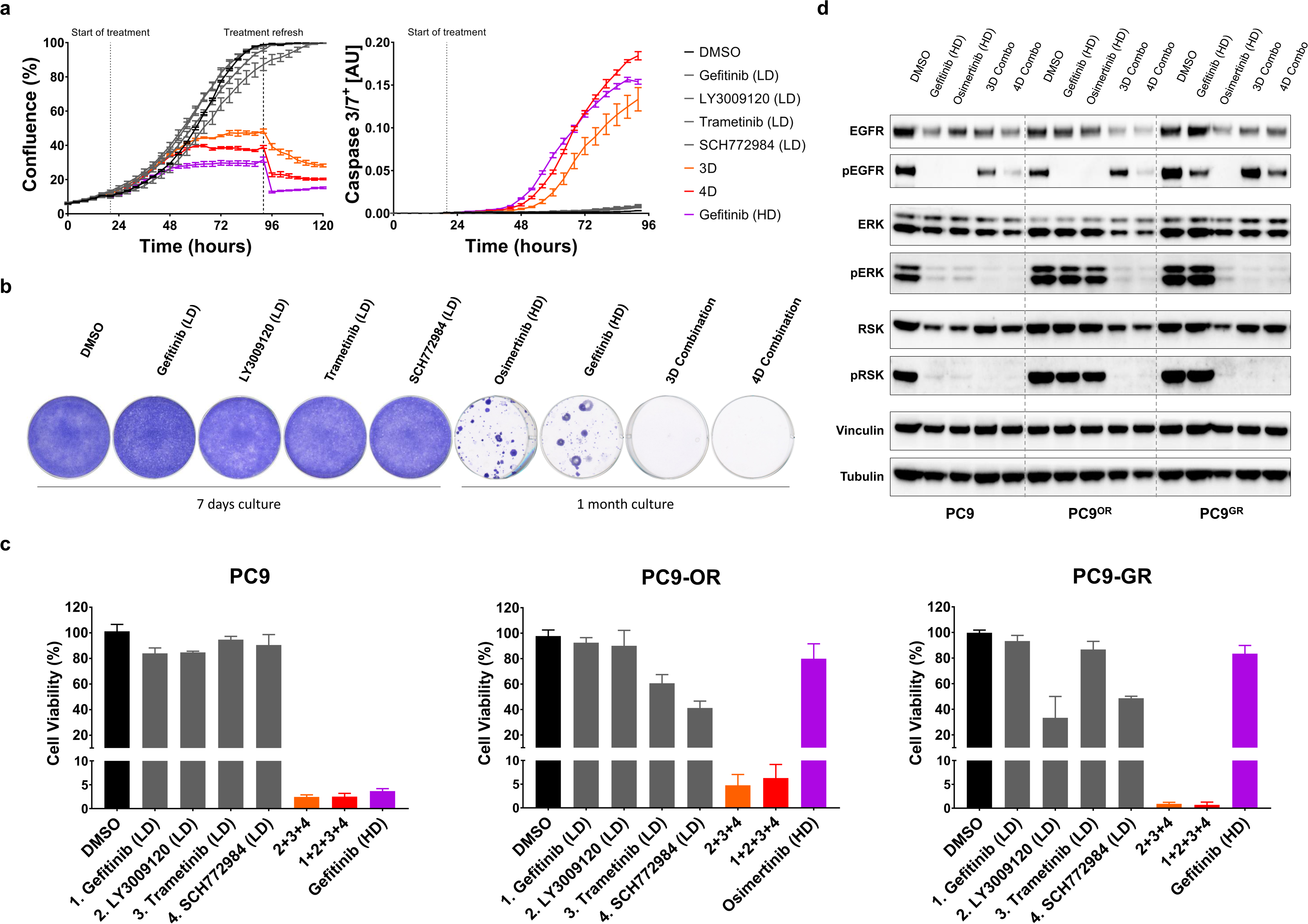
MLD therapy minimizes therapeutic resistance and is effective in EGFRi-resistant PC9 cells. **a,** MLD therapy abrogates cell proliferation and induces apoptosis in PC9 cells. PC9 cells were plated and incubated overnight to allow attachment to the plate. Cells were then treated with DMSO, with EGFR, RAF, MEK, ERK inhibitors at low dose, with 3D Combo (RAF+MEK+ERK inhibitors at LD) or with 4D Combo (EGFR+RAF+MEK+ERK inhibitors at LD) and placed in the IncuCyte^®^. Confluence (left) and caspase 3/7 activation (right) over time was measured by the IncuCyte^®^. Standard error of the mean (SEM) from 2 biologically independent replicates (each with 3 technical replicates) is plotted. **b,** MLD therapy prevents the acquisition of drug resistance in PC9 cells. PC9 cells were cultured with DMSO, with EGFR, RAF, MEK and ERK inhibitors at low dose (for 7 days) and with high dose (HD) of Osimertinib (200 nM), HD of Gefitinib (280 nM) and with 3D and 4D Combinations (for 1 month), after which plates were stained and scanned; A representative image from 3 biologically independent replicates is displayed. **c,** EGFRi-resistant PC9 cells remain sensitive to MLD therapy. PC9, PC9-OR (Osimertinib-resistant) and PC9-GR (Gefitinib-resistant) cells (see methods) were cultured with DMSO, with low doses of EGFR, RAF, MEK or ERK inhibitors, with 3D or 4D combinations or with HD of Gefitinib or Osimertinib for 4 days, after which cell viability was measured using CellTiter-Blue^®^. Standard deviation (SD) from 3 biologically independent replicates (each with 3 technical replicates) is plotted. **d,** MLD therapy blocks MAPK pathway in EGFRi-resistant PC9 cells. PC9, PC9-OR and PC9-GR cells were cultured with DMSO, HD of Osimertinib, HD of Gefitinib or with 3D or 4D combinations. Protein for western blotting was harvested after 24 hours of treatment; The level of pathway inhibition was measured by examining pERK and pRSK protein levels and the level of EGFR inhibition was measured by examining pEGFR protein levels in the western blot. Tubulin and Vinculin were used as loading control. A representative image from 2 biological replicates is displayed.

To study further if EGFRi-resistant cells are indeed sensitive to 3D and 4D combinations, we generated PC9 cells resistant to clinically-used EGFR inhibitors. We cultured PC9 cells in the presence of gefitinib (PC9-GR) or osimertinib (PC9-OR) until cells were no longer responsive to the inhibitors (see methods). We then tested the sensitivity of the resistant lines to 3D and 4D combinations. In both resistant cell populations, we saw an almost complete inhibition of cell viability after only 4 days of MLD therapy treatment and a complete MAPK pathway signalling blockade (Fig. 2c, d).

We then tested if the MLD strategy would also be effective in additional *in vitro* tumour models. After low dose determination (Supplemental Figs. 3a-c and Supplemental Table 1) we tested the MLD strategy in patient-derived (colorectal and NSCLC) organoids. Treatment with 3D and 4D combinations resulted in a major reduction in cell viability (Fig. 3a). In addition, we tested 6 different MAPK pathway addicted cell lines: HCC827 and H3255 (*EGFR* mutant lung cancer), H2228 and H3122 (*EML4-ALK* translocated lung cancer, in which EGFRi was replaced with ALK inhibitor crizotinib in the 4D combination), DiFi and Lim1215 (EGFR dependent colorectal cancer) and in 2 different PI3K pathway addicted cell lines: SKBR3 and HCC1954 (*HER2* amplified breast cancer, in which 4D combination consisted of HER2, PI3K, AKT and mTOR inhibitors). When treated with 4D combination, proliferation of all cell lines was inhibited, regardless of the tumour type/driver/genotype, pointing towards a broad applicability of the MLD treatment strategy (Supplemental Fig. 3d).

**Figure 3:**
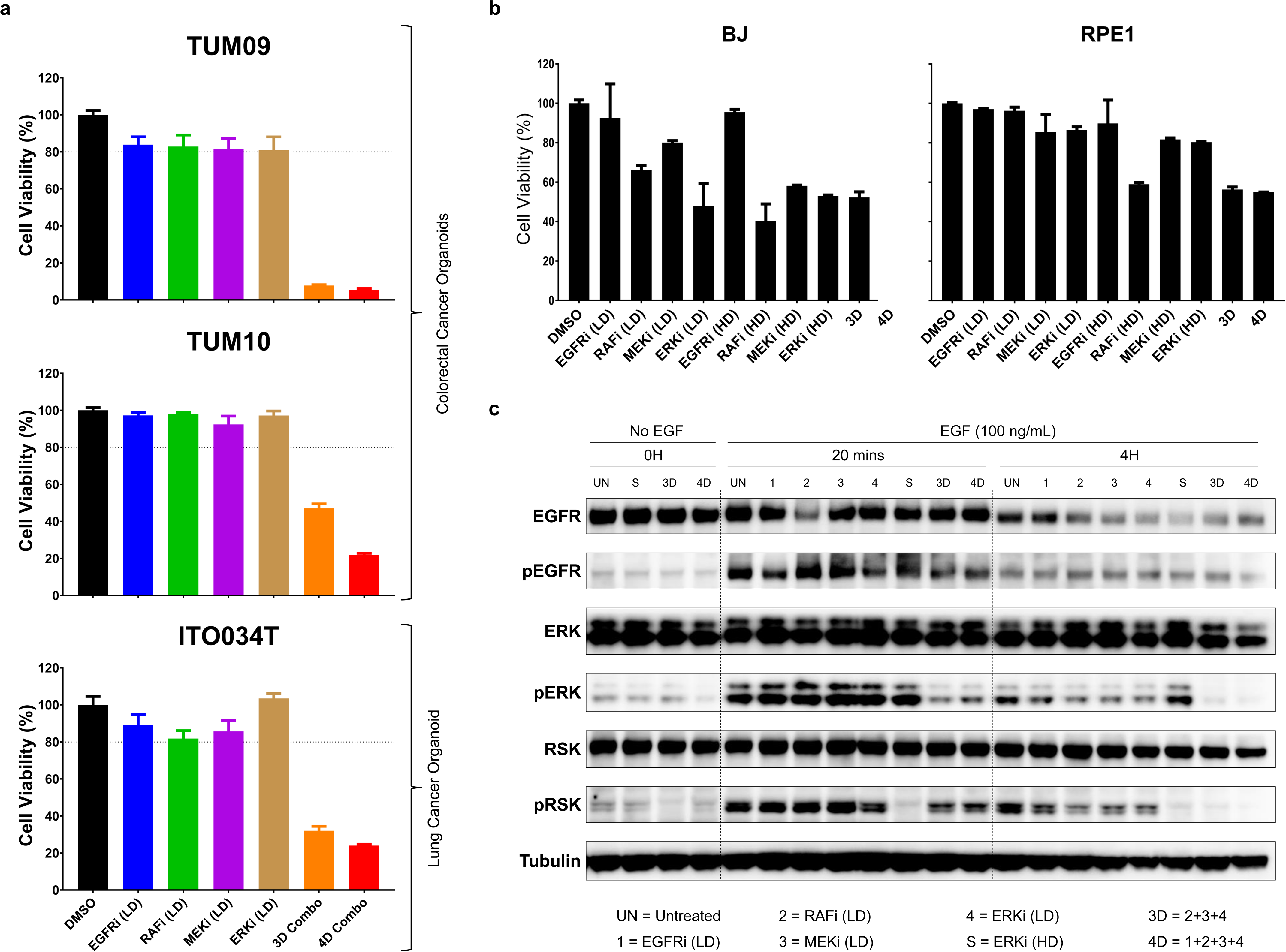
MLD therapy is effective in patient-derived organoids and is tolerated by normal cell lines. **a,** MLD therapy is effective in several colorectal and lung cancer patient-derived organoids. Organoids were cultured with DMSO, with EGFR, RAF, MEK and ERK inhibitors at LD and with 3D and 4D combos. After 5 days of drug treatment cell viability was measured using CellTiter-Glo^®^. SEM from 3 biologically independent replicates (each with 3 technical replicates) is plotted. **b,** Cell viability of normal cells is much less affected by MLD therapy than tumour cells. BJ and RPE1 cells were treated with DMSO, with EGFR, RAF, MEK and ERK inhibitors at low and high doses and with 3D and 4D Combos (using the LD and HD concentrations determined for PC9 cells). After 4 days of drug treatment cell viability was measured. SD from 3 replicates is plotted. **c,** MLD therapy allows pulsed signaling in normal cells. BJ cells, after overnight starvation, were treated with the indicated inhibitors/concentrations for 2 hours, after which EGF (100 ng/mL) was added. Cells were harvested before, 20 minutes and 4 hours after EGF stimulation. The level of pathway inhibition was measured by examining pERK and pRSK protein levels. The level of EGFR inhibition was measured by examining pEGFR protein levels in the western blot. Tubulin was used as loading control. A representative image from 2 biological replicates is displayed.

One of the major concerns when using multiple drugs in combination is the possible toxicity to normal tissues^24^. To address this, we tested the effect of the MLD strategy in primary human BJ (fibroblasts) and RPE1 (retinal pigment epithelium) cells. Upon 3D and 4D MLD drug combination treatment, cell viability was reduced but to a much lesser extent than in cancer cells, indicating that the MLD strategy might be tolerated by normal tissues (Fig. 3b). Since the MAPK pathway is rich in cross-talk and feedback control circuits^25, 26^, we also tested how a pulse of signalling through the EGFR pathway would be affected by 3D or 4D MLD treatment. We serum-starved BJ cells overnight and then incubated with 3D or 4D MLD drug combinations for two hours. After this, cells were stimulated with 100 ng/ml of EGF in the presence of 3D or 4D drug combinations. Twenty minutes after EGF stimulation, a significant amount of p-RSK was detected, which was no longer detected at 4 hours post EGF stimulation (Fig. 4c). These data suggest that the efficient inhibition of MAPK signalling exerted by 3D and 4D MLD treatment is the result of an effect of these drugs on homeostatic feedback/cross-talk signalling^25, 27^, as pulsatile signalling through the MAPK pathway seems to be much less affected than persistent signalling through an oncogene-activated MAPK pathway.

**Figure 4:**
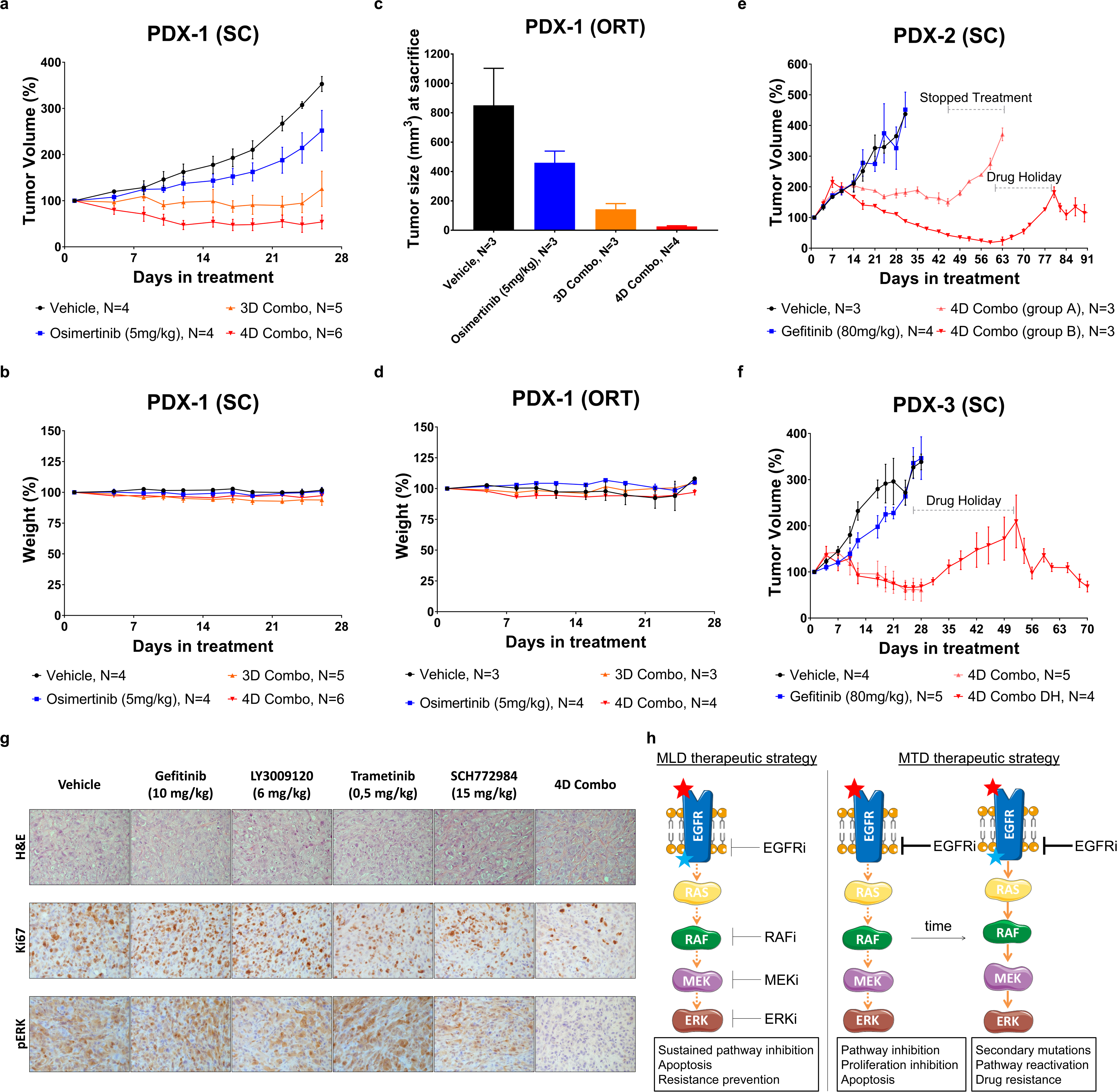
MLD therapy induces tumour regression without toxicity *in vivo*. **a-f,** Patient derived xenografts (PDX) are sensitive to MLD therapy. PDX tumours (see Supplemental Table 2) were implanted subcutaneously (a, e, f) or orthotopically in the lungs (c) of Crl:NU-Foxn1nu mice. PDX-1 was derived from a biopsy of a patient who became resistant to 3rd generation EGFR inhibitor osimertinib. PDX-2 and PDX-3 were derived from biopsies of patients who became resistant to 1st generation EGFR inhibitor erlotinib. PDX1 was implanted both subcutaneously (a) and orthotopically in the lungs (c). We defined the *in vivo* LD as 20-30% of the MTD for each of the individual drugs - gefitinib (10 mg/kg), LY3009120 (6 mg/kg), trametinib (0.5 mg/kg) and SCH772984 (15 mg/kg). After tumour establishment, mice were treated 5 days/week with vehicle, with osimertinib (5 mg/kg) and with 3D or with 4D Combos for 26 days, after which mice were sacrificed. In (a) tumour volume percentages ± SEM is shown, in (c) tumour size (mm^3^) at sacrifice ± SEM is shown and in (b) and (c) the mice weight percentages ± SEM is shown. (e) After tumour establishment, mice were treated 5 days/week with Vehicle (N=3), with gefitinib (80 mg/kg) (N=4) or with 4D Combo for 6 weeks (group A, N=3) or with 4D Combo for 8 weeks (group B, N=3). Mice treated with vehicle and gefitinib were sacrificed when tumours reached ∼2000mm3. After 6 weeks, Group A was taken off treatment and mice were sacrificed when tumours reached ∼2000mm3. After 8 weeks Group B was taken off treatment and was given 3 weeks of drug holiday. Mice were then treated for another 2 weeks with 4D combo, after which they were sacrificed. Tumour volume percentages ± SEM is shown. (e) After tumour establishment, mice were treated 5 days/week with vehicle (N=4), with gefitinib (80 mg/kg) (N=5) or with 4D Combo (N=9) for 4 weeks, after which mice were sacrificed, except for 4 animals from the 4D Combo group. These 4 mice were spared and were given 3 weeks drug holiday (4D Combo DH group), followed by another 3 weeks of treatment, after which they were sacrificed. Tumour volume percentages ± SEM is shown. **g,** Representative H&E, Ki67 and pERK stainings from tumour sections of PC9 xenografts are displayed. **h,** Schematic representation of the MLD therapy for the treatment of EGFR mutant NSCLC. While MTD therapy often results in secondary mutations which ultimately lead to resistance, MLD therapy is able to minimize therapeutic resistance even in EGFRi-resistant tumours without toxicity, both *in vitro* and *in vivo*.

To address if the MLD strategy is effective *in vivo*, we used patient derived xenograft (PDX) tumours from four different patients who had developed resistance to first-or second-line therapy with EGFR inhibitors erlotinib or osimertinib^6^ in the clinic by acquiring EGFR T790M mutation, *KRAS* mutation or *MET* amplification (Supplemental Table 2). For the *in vivo* studies we defined LD as 20-30% of the published maximum tolerated dose (MTD) for each of the individual drugs^18, 19, 28, 29^. Osimertinib-resistant PDX-1 was implanted subcutaneously and orthotopically in the lungs. In both models, treatment with 3D or 4D combination resulted in a reduction in tumour volume, without associated toxicity (Fig. 4a-d). Interestingly however, treatment with 4D combo was slightly more effective than 3D combo. Due to this finding we focused the *in vivo* studies that followed on the 4D combination. In all PDX models tested we observed similar results to PDX-1, i.e., a reduction in tumour volume, without significant toxicity (Figs. 4e, f and Supplemental Fig. 4e). Additionally, in gefitinib-resistant models PDX-2 and PDX-3 we tested if it would be possible to acquire resistance to the 4D MLD combination therapy during a drug holiday. In both PDX models, re-starting of 4D MLD therapy after a drug holiday resulted in a second response to the drug combination, indicating that overt resistance had not developed *in vivo* (Figs. 4e, f).

We also implanted PC9 cells in nude mice and treated them with vehicle, with EGFR, RAF, MEK and ERK inhibitors individually at low dose and with 4D combination. The use of low dose regimens was inadequate to suppress PC9 tumour growth when used as single agents, but when used in combination we observed a sustained reduction in the tumour volume of PC9 xenografts over a period of 70 days, which was associated with an extended survival (Supplemental Fig. 4a, b). These observations are also supported by immunohistochemical staining of the tumours, which show decreased Ki67 (a proliferation marker) and pERK (MAPK activation) levels in the tumours treated with 4D combination (Fig. 4g). Significantly, mice treated with 4D combination did not show any significant signs of toxicity, assessed by the weight of the mice over time and by the morphology of the GI tract and bone marrow (Supplemental Figs. 4c, f). In the clinic, the T790M mutation is already present (at very low percentages) in the majority of the tumours before undergoing anti-EGFR treatment^30, 31^. To mimic this scenario, we implanted in nude mice a mix of PC9 cells and PC9^GR^ cells (which are T790M positive) in a 9:1 ratio, respectively. Mice were treated with vehicle, with MTD of gefitinib and with 4D combination. Treatment with MTD of gefitinib resulted in a quick reduction of tumour volume which was followed by outgrowth of resistant cells, unlike the mice treated with 4D combination, where a sustained tumour control was observed (Supplemental Fig. 4d).

Despite the significant tumour regressions observed in the *in vivo* experiments none of the mice were fully cured, unlike in the *in vitro* data where all the cells were killed by the 3D or 4D combinations. To study why this is the case we studied the pharmacokinetics and pharmacodynamics of the four drugs *in vivo* over time. We found that drug plasma concentrations of gefitinib and trametinib dropped relatively slowly (T_1/2_ 8 hours), but the pan-RAF and ERK inhibitors were less stable in plasma (T_1/2_ of 5 and 4 hours, respectively). A similar difference was seen for intra-tumoural drug concentrations (Supplemental Figs. 5a, b). Consistent with this, we observed a complete inhibition of pRSK in tumour biopsies two hours after 4D combination drug administration, which progressively decreased after 8 and 24 hours (Supplemental Fig. 5c). These data indicate that, unlike in the *in vitro* experiments, two of the four drugs were not present at a significant concentration during at least 12 hours of the 24-hour treatment cycle. As a result of this, a sustained MAPK pathway inhibition was not achieved *in vivo*, possibly explaining why we didn’t achieve full tumour regressions. We tested this hypothesis *in vitro*, by removing RAF and ERK inhibitors from the treatment for approximately 8 hours every day. We found that, as hypothesized, when the drugs in the 3D or 4D combination were not present continuously the MLD therapy became less effective (Supplemental Fig. 5d).

We report here that treatment of *EGFR* mutant NSCLCs with MLD therapy effectively suppresses development of drug resistance, without associated toxicity. As such, our data challenge the paradigm that patients should be treated with the MTD of a targeted agent. Our data are consistent with a model in which diffuse inhibition of an oncogenic pathway at multiple nodes reduces selective pressure on each of the nodes to mutate and thereby increase response time (Fig 5i). Our findings also challenge the current model for MAPK pathway signalling, which postulates that the MAPK kinase cascade functions to amplify signals. Such amplification cascade model is clearly at odds with the data obtained here in which a very partial inhibition of each of 4 nodes in this cascade adds up to complete pathway inhibition. Further mechanistic studies are required to better understand the efficacy of the MLD strategy.

Importantly, we show that tumours having the most common mechanisms of clinically-observed resistance to high dose standard of care EGFR inhibitors still respond to MLD therapy. Therefore, MLD treatment strategy appears especially promising for patients that have already developed resistance to all clinically used EGFR inhibitors, including osimertinib. In such resistant tumours, multiple metastases may be present having different resistance mutations. In this study we have shown that MLD therapy is effective in PDX models having diverse EGFR inhibitor resistance mechanisms, including EGFR T790M mutation, *KRAS* mutation and *MET* amplification, highlighting that MLD therapy could apply to a diverse range of EGFR TKI resistant tumours. Indeed, in clinical practice, an MLD treatment strategy can only be tested in patients having developed resistance to standard-of-care EGFR inhibitors. We find in PDX models that 4D MLD is consistently somewhat better than 3D MLD, which may relate to the notion that not all *EGFR* alleles in the tumour may have acquired resistance mutations to the EGFR inhibitor therapy. Furthermore, it will be important to maintain osimertinib in an experimental MLD therapy trial, as this drug crosses the blood-brain barrier, and such late-stage patients may have (latent) brain metastases. We therefore suggest that clinical testing of the MLD strategy should include osimertinib.

While we never observed development of resistance to MLD therapy *in vivo*, even after long drug exposure, we did not achieve complete tumour regressions. This is most likely due to the short half-lives of the RAF and ERK inhibitors used in this study, which resulted in a situation in which we did not achieve a continuous pathway blockade. This may be improved by using continuous release formulations of these drugs, or by using drugs with longer half-lives.

The MLD therapy described here is fundamentally different from metronomic chemotherapy^32, 33^. In this latter scenario, low doses of chemotherapy are given at high frequency with the aim to suppress division of endothelial cells of the tumour vasculature. In the present MLD schedule, we target the MAPK pathway of the tumour itself, as growth inhibition in all cases parallels inhibition of the MAPK pathway (as judged by pRSK). Three-drug combinations given at MTD have been used before in pre-clinical^14^ and clinical studies^12, 13^ for *BRAF^V600E^*mutant tumours, showing clear therapeutic benefits, but such regimen have an associated cost of toxicity.

The lack of significant toxicity of the MLD therapy may be explained by the fundamentally different nature of MAPK pathway signalling between normal and EGFR mutant cancer cells. In the former, signalling is transiently activated when growth factors are present. In the latter, oncogenic mutations result in persistent activation of the pathway. Importantly, we show here that transient signalling in normal cells is, at least initially, not interrupted by MLD treatment (Fig. 3c). This may explain why long-term exposure of mice to MLD treatment is without major toxicity, as judged by lack of weight loss and lack of toxicity to gut epithelium and bone marrow.

Even though we focused mostly on EGFR mutant NSCLC, we have also shown that the MLD strategy can potentially be effective in other tumour types. Overall, our findings challenge the current paradigm of using the maximum tolerated dose of single targeted cancer drugs and suggest that, instead, it might be more beneficial to use a combination of multiple drugs that target the oncogenically activated pathway using sub-optimal drug concentrations.

## Acknowledgements

We thank A. Bardelli (Torino, Italy) for gift of cell lines and the facilities of The Netherlands Cancer Institute: Mouse Clinic – Intervention Unice (Natalie Proost, Bjorn Siteur, Bas van Gerwen, Charlotte Baak, Renske Grimmerink, Rebecca Theeuwsen and Marieke van de Ven), Robotics and Screening Center (Ben Morris and Roderick Beijersbergen), Clinical Pharmacology (Levi Buil and Artur Burylo), Experimental Animal Pathology, Flow Cytometry and Sequencing. We also thank Richard Marais for discussions. This work was supported by a grant from the Dutch Cancer Society through the Oncode Institute.

Alberto Villanueva was supported by the Fondo de Investigaciones Sanitarias, FIS (PI16-01898 (to A.V.), and by the Spanish Association Against Cancer, AECC (CGB14142035THOM) and Ideas Semilla project (IDEAS098VILL-IDEAS16) and Generalitat de Catalunya (2014SGR364) (to A.V.). L.F received a European Union’s Horizon 2020 research and innovation programme under the Marie Sklodowska-Curie, grant agreement number 799850 (L.F). E.N. received support from the SLT006/17/00127 grant, funded by the Department of Health of the Generalitat de Catalunya by the call “Acció instrumental d’intensificació de professionals de la salut” and the AECC grant from Junta Provincial de Barcelona. We thank CERCA Program/Generalitat de Catalunya for their institutional support and grant 2017SGR448.

## Authors’ Contributions

R.B. supervised all research. J.N., A.V., O.vT, and R.B., wrote the manuscript. J.N., E.N., S.O., E.B., L.F., C.M., S.K., A.J., H.H., L.W., designed, performed and analysed the experiments. E.V., A.V., E.F., A.M-M and Lo.W. provided advice for the project. All authors commented on the manuscript.

## METHODS

### Cell lines culture and drug-response assays

The PC9 cell line was obtained from ATCC. PC9^OR^ (osimertinib-resistant) cells were made by continuous (2 months) drug exposure of PC9 cells to osimertinib (AZD9291). PC9^GR^ (gefitinib-resistant) cells were made by continuous (2 months) drug exposure of PC9 cells to gefitinib - these cells were sequenced and tested positive for the EGFR T790M mutation. The HCC827, H3255, H3122, H2228, SKBR3, HCC1954, BJ and RPE1 cell lines were obtained from ATCC. And DiFi and Lim1215 cell lines were a gift from A. Bardelli (Torino, Italy). BJ and RPE1 cells were cultured in DMEM (Gibco 41966029). SKBR3 and HCC1954 which were cultured in DMEM/F-12 medium (Gibco 31331028). All the other cell lines were cultured in RPMI medium (Gibco 21875034). All the cell lines media were supplemented with 10% FBS (Serana), 1% penicillin/streptomycin (Gibco 15140122) and 2 mM L-glutamine (Gibco 25030024). All cell lines were cultured at 37°C and with 5% CO_2_. All cell lines were validated by STR profiling and mycoplasma tests were performed every 2-3 months.

All drug-response assays were performed in triplicate, using black-walled 384-well plates (Greiner 781091). Cells were plated at the optimal seeding density (Supplemental Table 1) and incubated for approximately 24 hours to allow attachment to the plate. Drugs were then added to the plates using the Tecan D300e digital dispenser. 10 µM phenylarsine oxide was used as positive control (0% cell viability) and DMSO was used as negative control (100% cell viability). Four days later, culture medium was removed and CellTiter-Blue (Promega G8081) was added to the plates. After 1-4 hours incubation, measurements were performed according to manufacturer’s instructions using the EnVision (Perkin Elmer).

### Organoid culture and drug-response assays

Colorectal (CRC) and non-small cell lung cancer (NSCLC) organoids were established and handled as previously described^1^. All drug-response assays were performed in replicate, each by independent researchers. PDOs were mechanically and enzymatically dissociated into single cells, pipetted through a 40 µM cell strainer, and re-plated to allow for organoids formation. At day 4 PDOs were collected, Cultrex was removed by incubation of the cell pellet in 1 mg/ml dispase II (Sigma D4693) for 15 minutes. Whole organoids were counted using a hemocytometer and trypan blue. PDOs were resuspended in 1:3 Advanced Dulbecco’s Modified Eagles Medium with Nutrient Mixture F-12 Hams (Ad-DF) (Invitrogen 12634), supplemented with 1% penicillin/streptomycin (Invitrogen 15140122), 1% HEPES (Invitrogen 15630056) and 1% GlutaMAX (Invitrogen 35050) (Ad-DF+++):Cultrex at a concentration of 20 organoids/µl. Five µl/well was dispensed in clear-bottomed, white-walled 96-well plates (Greiner Bio-One 655098) and overlaid with 200 µl CRC or NSCLC culture medium. We generated 10-step dose response curves using the Tecan D300e digital dispenser, interpolated IC_20_ values and re-screened organoids in presence of a range of concentration around the IC_20_ of each drug separately and in 3D and 4D Combos. In addition, we re-performed the dose-response curves to control for variation between experiments. Read-out was performed at day 10 in the positive control (10 µM phenylarsine oxide), negative control (DMSO), and the drug-treated wells. Quantification of cell viability was done by replacing the CRC medium with 50 µL Cell-TiterGlo 3D (Promega G9681) mixed with 50 µL Ad-F+++. Measurements were performed according to manufacturer’s instructions on an Infinite 200 Pro plate reader (Tecan Life Sciences) with an integration time of 100 ms.

### Compounds, reagents and antibodies

Gefitinib (100140), LY3009120 (206161), trametinib (201458), SCH772984 (406578), osimertinib (206426), crizotinib (202222), lapatinib (100946), BKM120 (204690), MK2206 (201913) and AZD8055 (200312) were purchased from MedKoo Biosciences. Erlotinib (S7786), BGB-283 (S7926), selumetinib (S1008) and LY-3214996 (S8534) were purchased from Selleckchem. Annexin V-FITC Apoptosis Staining Detection Kit was purchased from Abcam (ab14085).

Antibodies against Tubulin (T9026) and Vinculin (V9131) were purchased from Sigma; antibodies against EGFR (4267), pERK (4377), ERK (9102) and RSK (8408) were purchased from Cell Signalling; antibody against pRSK (04-419) was purchased from Millipore; antibody against pEGFR (ab5644) was purchased from Abcam.

### Colony formation and IncuCyte cell proliferation assays

Cells were seeded in the appropriate density (Supplemental Table 1) in 6-well plates. Cells were incubated for approximately 24 hours to allow attachment to the plates, after which drugs were added to the cells using the Tecan D300e digital dispenser as indicated. The culture media/drugs were refreshed every 3/4 days. When control wells (DMSO) were confluent (unless otherwise stated in the text) cells were fixed using a solution of 2% formaldehyde (Millipore 104002) diluted in phosphate-buffered saline (PBS). Two hours later, they were stained, using a solution of 0.1% crystal violet (Sigma HT90132) diluted in water. Not more than 10 minutes later the staining solution was removed, plates were washed with water left to dry overnight. Finally, plates were scanned and stored.

For IncuCyte proliferation assays, cells were seeded in 96-well plates and incubated overnight to allow attachment to the plates. Drugs were added to the cells using the Tecan D300e digital dispenser. Cells were imaged every 4 hours in the IncuCyte ZOOM (Essen Bioscience). Phase-contrast images were collected and analysed to detect cell proliferation based on cell confluence. For cell apoptosis, IncuCyte® Caspase-3/7 green apoptosis assay reagent (Essen Bioscience 4440) was also added to culture medium and cell apoptosis was analysed based on green fluorescent staining of apoptotic cells.

### Western Blots

After the indicated culture period, cells were washed with chilled PBS and then lysed with RIPA buffer (25mM Tris - HCl pH 7.6, 150mM NaCl, 1% NP-40, 1% sodium deoxycholate, 0.1% SDS) containing protease inhibitors (Complete (Roche) and phosphatase inhibitor cocktails II and III). Samples were then centrifuged for 10 minutes at 14.000 rpm at 4°C and supernatant was collected. Protein concentration of the samples was normalized after performing a Bicinchoninic Acid (BCA) assay (Pierce BCA, Thermo Scientific), according to the manufacturer’s instructions.

Protein samples (denatured with DTT followed by 5 minutes heating at 95°C) were then loaded in a 4-12% polyacrylamide gel. Gels were run (SDS-PAGE) for approximately 60 minutes at 165 volts. Proteins were then transferred from the gel to a polyvinylidene fluoride (PVDF) membrane, using 330 mA for 90 minutes.

After the transfer, membranes were placed in blocking solution (5% bovine serum albumin (BSA) in PBS with 0,1% Tween-20 (PBS-T). Subsequently, membranes were probed with primary antibody in blocking solution (1:1000) and left shaking overnight at 4°C. Membranes were then washed 3 times for 10 minutes with PBS-T, followed by one hour incubation at room temperature with the secondary antibody (HRP conjugated, 1:10000) in blocking solution. Membranes were again washed 3 times for 10 minutes in PBS-T. Finally, a chemiluminescence substrate (ECL, Bio-Rad) was added to the membranes and the Western Blot was resolved using the ChemiDoc (Bio-Rad).

### Mouse xenografts studies

All animal experiments were approved by the Animal Ethics Committee of the Netherlands Cancer Institute or by the Animal Ethics Committee of the Institut Català d’Oncologia and performed in accordance with institutional, national and European guidelines for Animal Care and Use.

PC9 cell line xenografts: One million PC9 cells were resuspended in PBS and mixed 1:1 with matrigel (Corning 354230). Cells were injected subcutaneously into the posterior flanks of 7-week-old immunodeficient BALB/cAnNRj-Foxn1nu mice (half male and half female; Janvier Laboratories, The Netherlands). Tumour formation was monitored twice a week. Tumour volume, based on calliper measurements, was calculated by the modified ellipsoidal formula (tumour volume = 1/2(length × width2)). When tumours reached a volume of approximately 200 mm^3^, mice were randomized into the indicated treatment arms. Vehicle, gefitinib, LY3009120, trametinib, SCH772984 or the combination of the 4 inhibitors were formulated in DMSO: Kolliphor EL (Sigma 27963): Saline solution, in a ratio of (1:1:8). Mice were treated 5 days a week (Monday to Friday) at the indicated doses by intraperitoneal injection.

Patient-derived xenografts (PDX) and orthotopic xenograft (PDOX): Primary tumours were obtained from Bellvitge Hospital (HUB) and the Catalan Institute of Oncology (ICO) with approval by the Ethical Committee. Ethical and legal protection guidelines of human subjects, including informed consent from the patient to implant the tumour in mice, were followed. PDX-1 was generated from a lung adenocarcinoma biopsy from a patient who was treated with Erlotinib (first line), Gefitinib + Capmatinib (second line) and Cisplatin+Pemetrexed (third line). This tumour has an EGFR mutation (del19) and *MET* amplification. PDX-2 was generated from a lung adenocarcinoma biopsy from a patient who was treated with Erlotinib (first line), Gefitinib + Capmatinib (second line) and Carboplatin+Gemcitabine and Nivolumab (third line). This tumour has an EGFR mutation (L858R) and *MET* amplification. PDX-3^2^ was generated from a lung adenocarcinoma biopsy of a brain metastasis from a patient who was treated with Erlotinib (first line) and Osimertinib (second line). PDX-4 was generated from a lung adenocarcinoma biopsy from a patient who was treated with Afatinib (first line) and CBDCA + pemetrexed (second line). This tumour has a germline p53 mutation and a EGFR mutation (del19). Tumours were isolated and implanted subcutaneously (or orthotopically, in the lungs, in the case of PDX-3) into Crl:NU-Foxn1nu mice by following previously reported procedures^2, 3^. In the subcutaneous models, tumour volume was monitored twice a week by a digital caliper. When tumours reached a volume of approximately 200-600 mm^3^, mice were randomized into the indicated treatment arms. In the orthotopic model, tumours were left to grow for 2 weeks, followed by 26 days of treatment. Vehicle, gefitinib, osimertinib or the 3D and 4D Combos were formulated in DMSO: Kolliphor EL (Sigma 27963): Saline solution, in a ratio of (1:1:8). Mice were treated 5 days a week (Monday to Friday) at the indicated doses by intraperitoneal injection.

### *In vivo* pharmacokinetics and pharmacodynamics studies

Plasma and tumour samples were assayed by liquid chromatography triple quadrupole mass spectrometry (LC-MS/MS) using an API4000 detector (Sciex) for the simultaneous determination of Gefitinib (MRM: 447.4/128.1), LY3009120 MRM: 425.5/324.2), Trametinib (616.3/491.2) and SCH772984 (MRM: 588.4/320.2). Gefitinib-d8 (MRM: 455.4/136.3) was used as internal standard. LC separation was achieved using a Zorbax Extend C18 column (100 × 2.0 mm; ID). Mobile phase A and B comprised 0.1% formic acid in water and methanol, respectively. The flow rate was 0.4 ml/min and a linear gradient from 20%B to 95%B in 2.5 min, followed by 95%B for 2 min, followed by re-equilibration at 20%B for 10 min was used for elution. Sample pre-treatment was accomplished by mixing 5 ul (plasma) or 25 ul (tumour homogenate) with 30 or 150 ul, respectively, of formic acid in acetonitrile (1+99) containing the internal standard. After centrifugation, the clear supernatant was diluted 1+4 with water and 50 ul was injected into the LC-MS/MS system.

The plasma/tumour samples were harvested at the time points indicated in supplemental figure 5. Blood samples were obtained by tail cut (at 2h and 8h time points) and by cardiac puncture at the 24h time point. Samples were collected on ice in tubes containing potassium EDTA as anticoagulant. The tubes were immediately cooled in melting ice and centrifuged (10 minutes, 5000g, 4°C) to separate the plasma fraction, which was transferred into clean vials. For the tumours samples, the mice were sacrificed by cervical dislocation, the tumour was dissected and frozen at −80°C. Half of the tumour was then lysed mechanically with RIPA buffer and lysates were analysed by Western blot. The other half was weighed and homogenized in 1 ml of ice-cold 1% of BSA in water and stored at −20°C until further analysis.

## SUPPLEMENTAL FIGURE LEGENDS

**Supplemental Figure 1:**
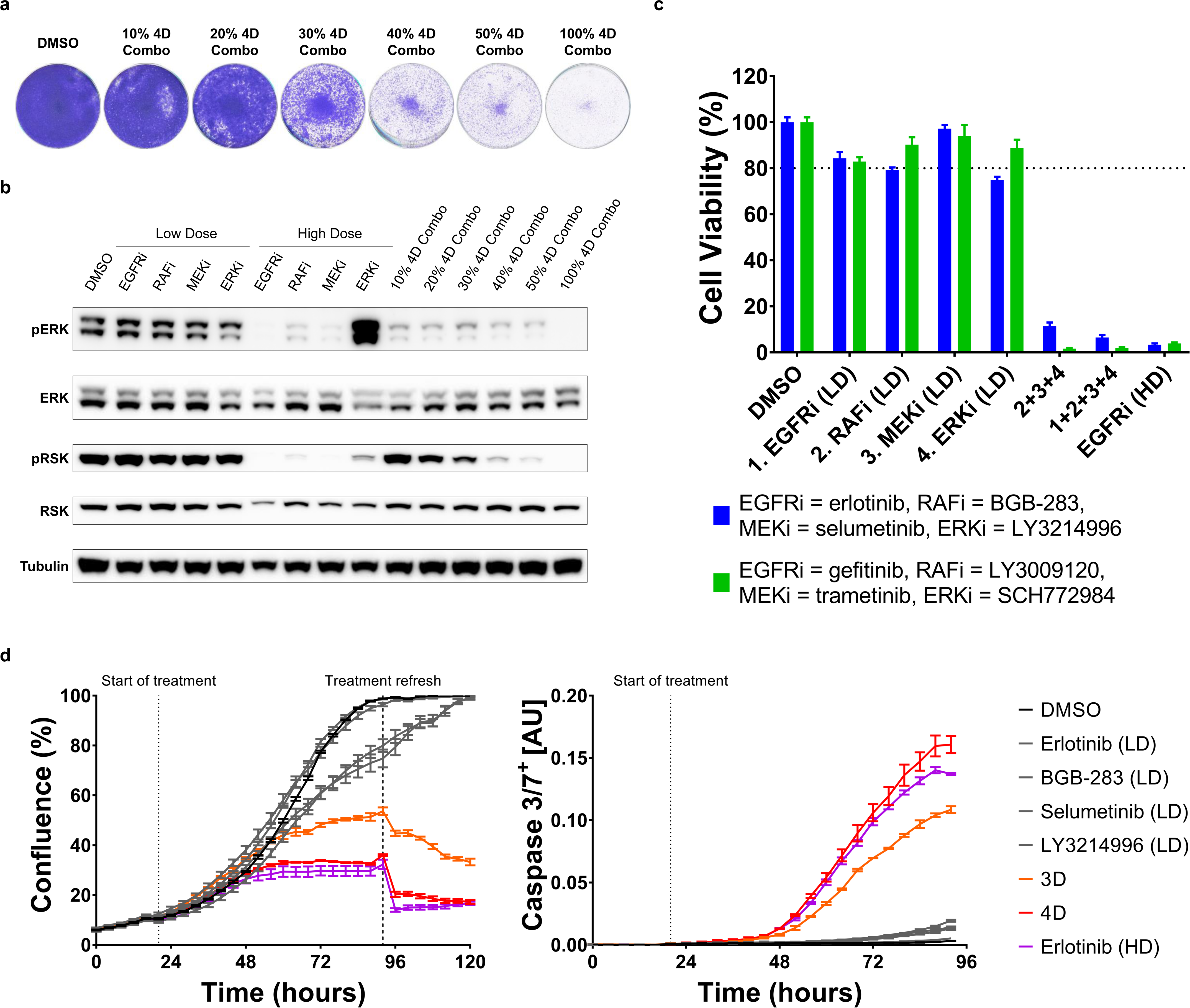
A drug concentration threshold is necessary for the efficacy of MLD therapy, which is not drug-specific. **a,** Dilution of 4D Combo results in incomplete inhibition of proliferation. PC9 cells were plated and incubated overnight to allow attachment to the plate. Cells were then treated with DMSO, with 4D Combo and with the indicated dilutions of 4D Combo. Cells were cultured for 7 days, after which plates were stained and scanned; A representative image from 3 biologically independent replicates is displayed. **b,** Dilution of 4D Combo results in incomplete MAPK pathway inhibition. PC9 cells were cultured with DMSO, with EGFR, RAF, MEK and ERK inhibitors both at low and at high doses, with 4D Combo and with different dilutions of 4D combo. Protein for western blotting was harvested after 48 hours of treatment. The level of pathway inhibition was measured by examining pERK and pRSK protein levels; Tubulin was used as loading control. A representative image from 2 biological replicates is displayed. **c, d,** MLD therapy efficacy is not drug-specific. PC9 cells were plated and incubated overnight to allow attachment to the plate; Cells were then treated with two different inhibitors for each of the nodes in the MAPK pathway (gefitinib or erlotinib as EGFRi, LY3009120 or BGB-283 as RAFi, Trametinib or selumetinib as MEK and SCH772984 or LY-3214996 as ERKi) as indicated. In (d) cell viability was measured using CellTiter-Blue® after 4 days of treatment. Standard deviation (SD) from 3 biologically independent replicates (each with 3 technical replicates) is plotted. In (e) the confluence (left) and caspase 3/7 activation (right) over time was measured by the IncuCyte®; 3 days after the first treatment the drugs and media were refreshed. SEM from 3 replicates is plotted.

**Supplemental Figure 2:**
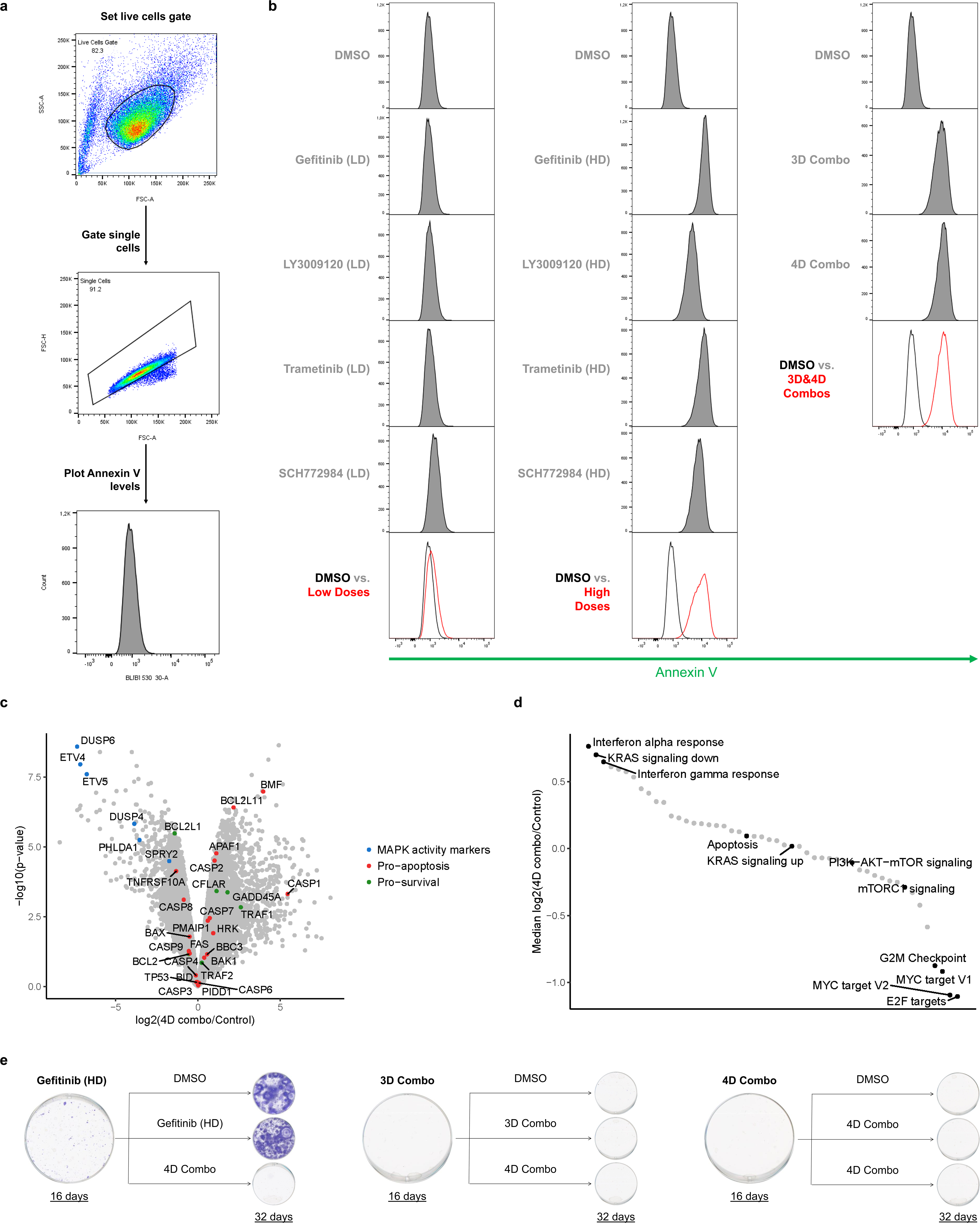
MLD therapy induces apoptosis and prevents drug resistance. **a,** Gating strategy used for (b). Live cells were gated from all events; then single cells were gated from the live cells and, finally, Annexin V levels were plotted from the single cells. **b,** 3D and 4D Combos induce apoptosis at comparable levels as high doses of each inhibitor in PC9 cells. PC9 cells were stained with Annexin V-FITC Apoptosis Staining/Detection kit (ab14085) after 48 hours of drug treatment. The Annexin V levels were measured by flow cytometry (BD LSRFortessa) and analysed using FlowJo 10. **c, d,** Transcriptome analysis of PC9 cells treated with 4D combo. (c) Volcano plot of differential gene expression analysis. (d) Median log2-fold change of the MSigDB hallmark gene-sets, ranked from high to low. For (c) and (d) PC9 cells were treated with DMSO for 48 hours or with 4D combo for 48 or 72 hours. Experiments were performed in duplicates. Because the difference between 48 and 72 hour 4D combo treatment was comparable to the variability between replicates, the four MLD treated samples were considered replicates. Differential expression analysis was performed using the R-package limma [Ritchie et al, 2015] and the MSigDB hallmark gene-sets analysis was performed using version 6.2 of MSigDB [Liberzon et al, 2015]. **e,** MLD therapy prevents the acquisition of drug resistance in PC9 cells. PC9 cells were cultured with high dose of gefitinib (280 nM) and with 3D and 4D Combos (4 plates per condition). After 16 days in culture, one plate was fixed and stained. From the remaining three plates (per condition) one was switched to DMSO treatment, the other was switched to 4D Combo and the third one continued with the previous treatment. Sixteen days later (after 32 days of “treatment” in total) cells were fixed and stained and then plates were scanned. A representative image from the 3 replicates performed is displayed.

**Supplemental Figure 3:**
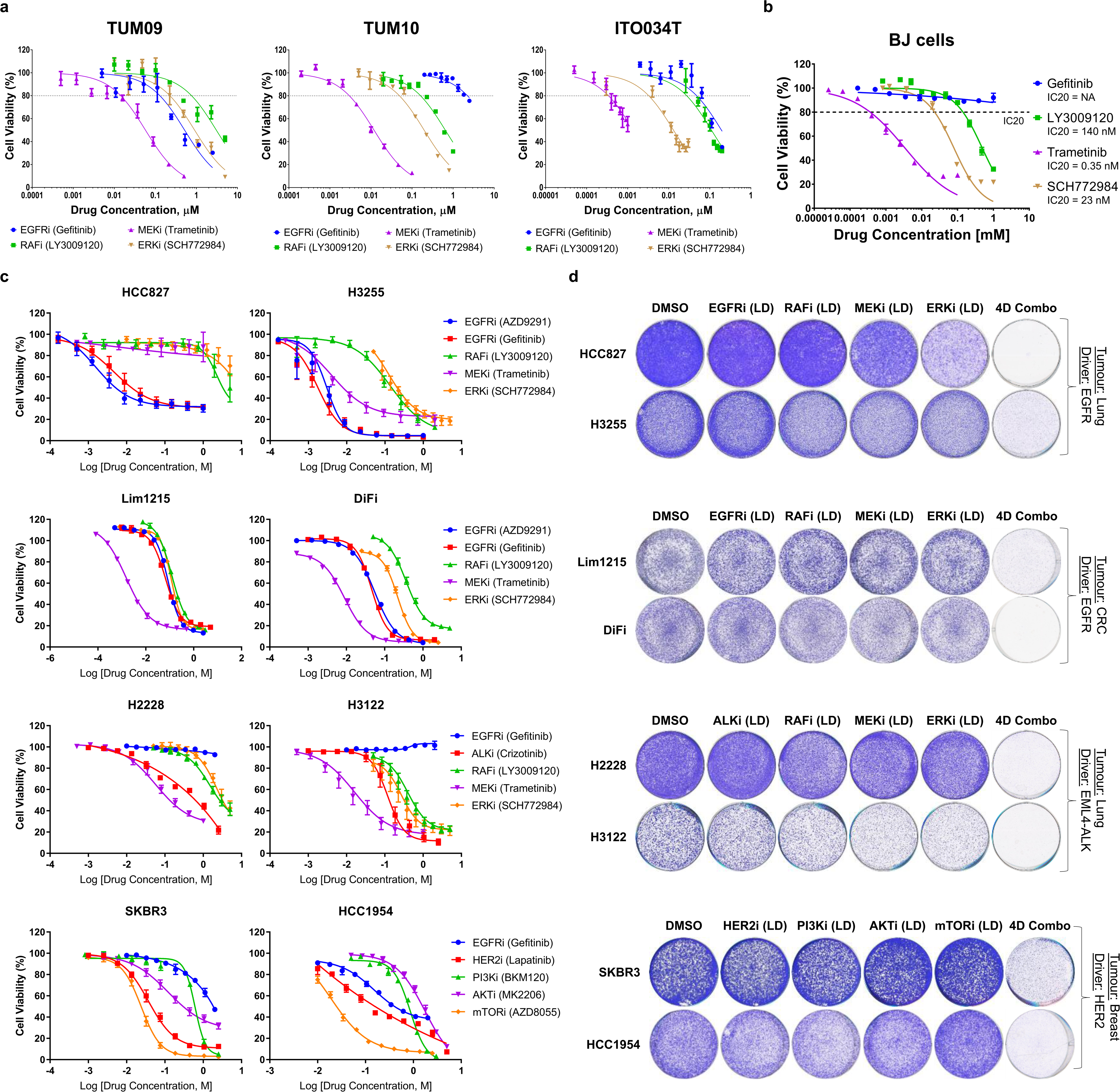
MLD therapy is effective in multiple cancer cell lines. **a-c,** Dose-response curves across the organoid and cell line panel. (a) Organoids were cultured with DMSO or with the different inhibitors and after 5 days of drug treatment cell viability was measured using CellTiter-Glo^®^. SEM from 3 biologically independent replicates (each with 3 technical replicates) is plotted. (b, c) Cells were plated in 384-well plates. Drugs were added ∼24h after plating; after 4 days of exposure to the drugs cell viability was measured using CellTiter-Blue^®^. SEM from 3 biologically independent replicates (each with 3 technical replicates) is plotted. Low doses (IC20s) were then determined (see Supplemental Table 1). **d,** MLD therapy is effective in several cell lines/tumour types. HCC827, H3255, Lim1215, DiFi, H2228, H3122, SKBR3 and HCC1954 cell lines were treated with DMSO, with the indicated pathway inhibitors at low dose and with their combination (4D Combo). After 10 days of treatment plates were stained and scanned; A representative image from 3 biologically independent replicates is displayed.

**Supplemental Figure 4:**
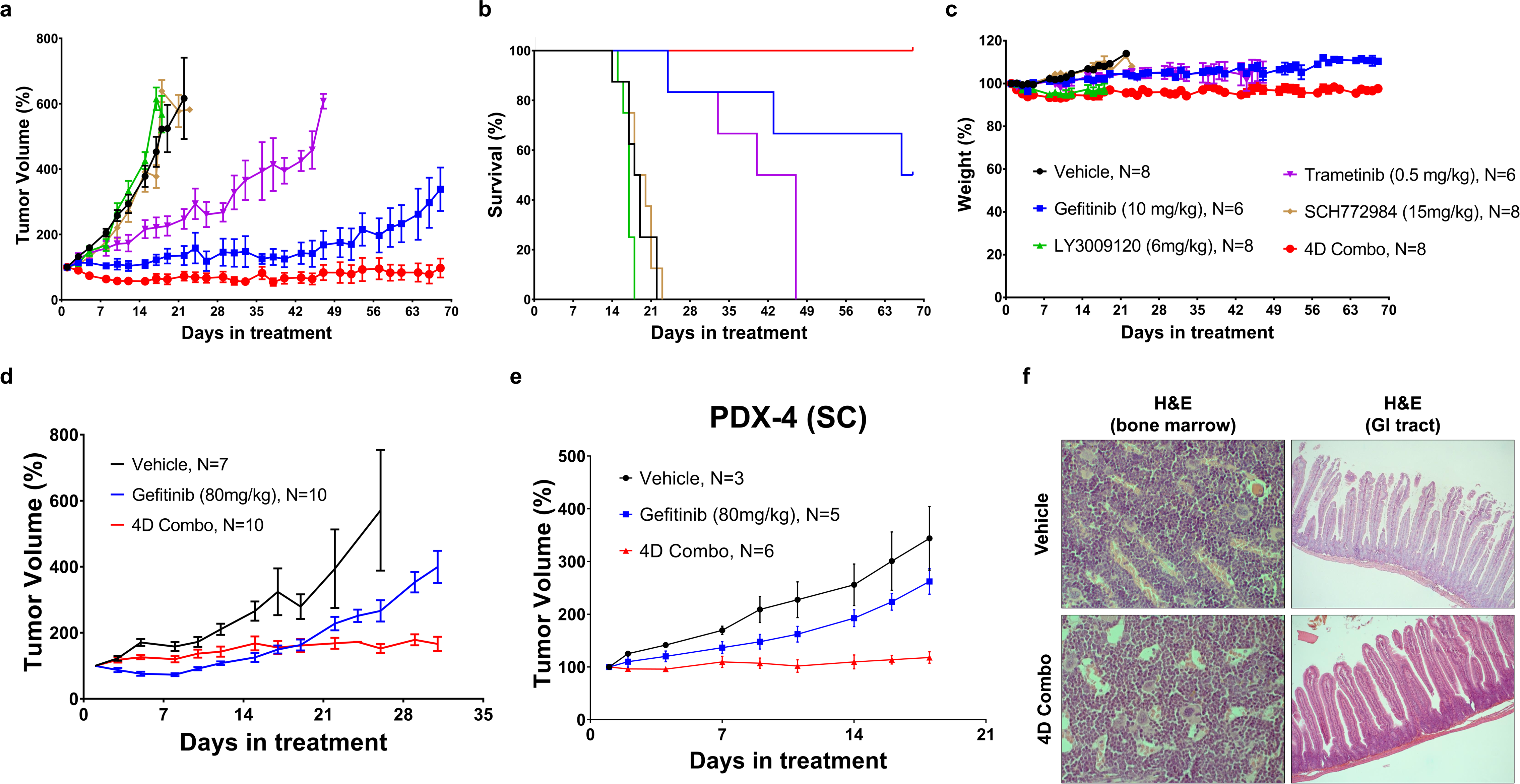
MLD therapy reduces tumour volume *in vivo* without toxicity. **a-d**, PC9 xenografts are sensitive to 4D Combo without toxicity. (a-c) PC9 cells were grown as tumour xenografts in BALB/cAnNRj-Foxn1nu mice. After tumour establishment (200–250 mm3), mice were treated 5 days/week with vehicle, gefitinib (10 mg/kg), LY3009120 (6 mg/kg), trametinib (0.5 mg/kg), SCH772984 (15 mg/kg) or the combination of the 4 inhibitors (4D Combo) for 10 weeks. In (a) the mean tumour volume percentages ± SEM is shown; In (b) the Kaplan-Meier survival curve is shown; In (c) the mice weight percentages ± SEM is shown. (d) PC9 cells and PC9^GR^ cells were mixed in a 9:1 ratio, respectively, and were grown as tumour xenografts in BALB/cAnNRj-Foxn1nu mice. After tumour establishment (200–250 mm3), mice were treated 5 days/week with vehicle, with the MTD of gefitinib (80 mg/kg) and with 4D Combo – cocktail containing gefitinib (1 mg/kg), LY3009120 (6 mg/kg), trametinib (0.1 mg/kg), SCH772984 (15 mg/kg) for 30 days. The mean tumour volume percentages ± SEM is shown. **e,** EGFR and p53 mutant PDX responds to 4D Combo. PDX-4 was generated from a biopsy of patient with EGFR and TP53 mutation that progressed after afatinib and chemotherapy treatment. After tumour establishment, mice were treated 5 days/week with Vehicle (N=3), with gefitinib (80 mg/kg) (N=5) or with 4D combo (N=6) – cocktail containing gefitinib (10 mg/kg), LY3009120 (6 mg/kg), trametinib (0.5 mg/kg) and SCH772984 (15 mg/kg) (N=6) for 18 days. Tumour volume percentages ± SEM is shown. **f,** Representative H&E stainings from the GI tract and the bone marrow of the PC9 xenografts in a-c are displayed.

**Supplemental Figure 5:**
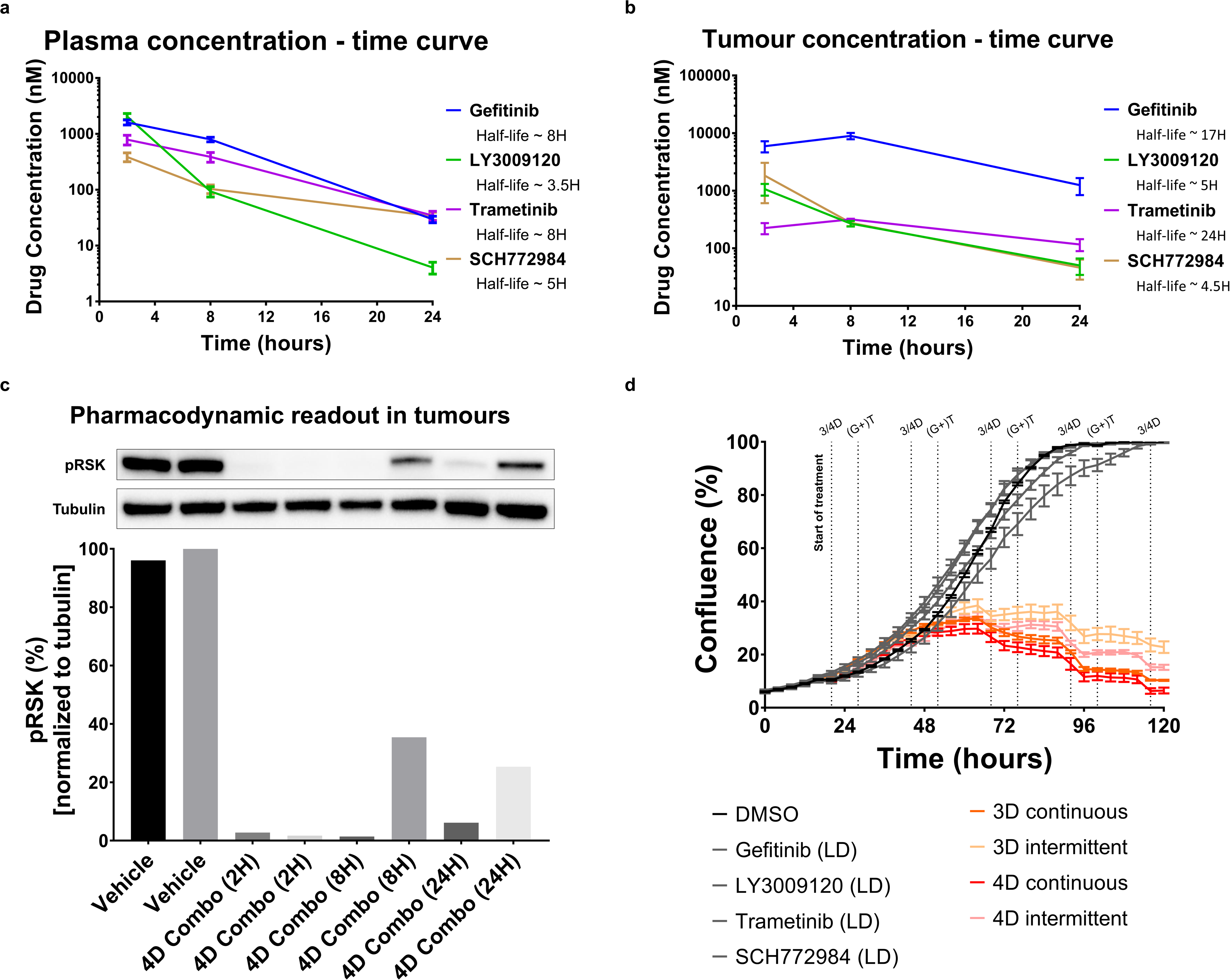
PK-PD studies in PC9 xenografts reveal different half-lives of the inhibitors. **a-c,** Pharmacokinetic and pharmacodynamics studies in PC9 xenografts. PC9 cells were injected (bilaterally) subcutaneously in BALB/cAnNRj-Foxn1nu mice. After tumour establishment (∼200 mm3), mice were treated with vehicle (N=4) or 4D Combo (N=12). Vehicle mice were sacrificed 2H after treatment; Mice treated with 4D combo were sacrificed 2, 8 and 24h after treatment, respectively; 4 mice were sacrificed per time point. Blood and tumours were harvested; half of the tumour was used for the PD study and the other half was used for biochemical analysis. The drug concentrations in the blood and in the tumours were determined by mass spectrometry. In (a) the concentration of the individual drugs in the plasma is displayed and in (b) the concentration of the individual drugs in the tumours is displayed; SEM is plotted. In (c) the level of pathway inhibition in the tumours was measured by examining pRSK protein levels in the western blot (WB). Tubulin was used as loading control. WB was quantified using the Image Lab software, from Bio Rad. **d,** Intermittent MLD therapy is less efficient in reducing cell growth in PC9 cells. PC9 cells were plated and incubated overnight to allow attachment to the plate. Cells were then treated with DMSO, with EGFR, RAF, MEK, ERK inhibitors at low dose, with 3D Combo or with 4D Combo. To mimic the availability of the drugs *in vivo*, in some of the 3D and 4D combo replicates the RAF and ERK inhibitors were removed from the culture media for approximately 8 hours every day (called intermittent MLD therapy). Confluence over time was measured by the IncuCyte^®^. SEM from 3 replicates is plotted.

**Supplemental Table 1:** Compendium of drivers, plating density, low doses and high doses for all the cell lines and organoids used in the study.

|  |  |  |  |  |  | Drug concentrations used in the study (µM) |  |  |  |  |  |  |  |
| --- | --- | --- | --- | --- | --- | --- | --- | --- | --- | --- | --- | --- | --- |
|  |  |  |  |  |  | Low Doses |  |  |  | High Doses |  |  |  |
| Cell Line Name | Tissue of Origin | Driver | Plating density (384-well plates) | Plating density (96-well plates) | Plating density (6-well plates) | EGFRi (Gefitinib) | RAFi (LY3009120) | MEKi (Trametinib) | ERKi (SCH772984) | EGFRi (Gefitinib) | RAFi (LY3009120) | MEKi (Trametinib) | ERKi (SCH772984) |
| PC9/PC9<sup>OR</sup>/PC9<sup>GR</sup> | Lung | EGFR | 150 cells | 2000 cells | 20000 cells | 0.007 | 0.25 | 0.008 | 0.25 | 0.28 | 2.8 | 0.32 | 2.5 |
| HCC827 |  | EGFR | 350 cells | NA | 40000 cells | 0.0025 | 0.5 | 0.1 | 0.75 | 1 | 5 | 1 | 5 |
| H3255 |  | EGFR | 750 cells | NA | 50000 cells | 0.001 | 0.03 | 0.001 | 0.04 | 1 | 2 | 2 | 5 |
| Cell Line Name | Tissue of Origin | Driver | Plating density (384-well plates) | Plating density (96-well plates) | Plating density (6-well plates) | ALKi (Crizotinib) | RAFi (LY3009120) | MEKi (Trametinib) | ERKi (SCH772984) | ALKi (Crizotinib) | RAFi (LY3009120) | MEKi (Trametinib) | ERKi (SCH772984) |
| H3122 | Lung | EML4-ALK | 1000 cells | NA | 60000 cells | 0.075 | 0.11 | 0.0038 | 0.084 | 2.5 | 5 | 1 | 5 |
| H2228 |  | EML4-ALK | 1500 cells | NA | 80000 cells | 0.046 | 0.5 | 0.01 | 0.75 | 2.5 | 5 | 1 | 5 |
| Cell Line Name | Tissue of Origin | Driver | Plating density (384-well plates) | Plating density (96-well plates) | Plating density (6-well plates) | EGFRi (Gefitinib) | RAFi (LY3009120) | MEKi (Trametinib) | ERKi (SCH772984) | EGFRi (Gefitinib) | RAFi (LY3009120) | MEKi (Trametinib) | ERKi (SCH772984) |
| DiFi | CRC | EGFR | 1000 cells | NA | 60000 cells | 0.02 | 0.17 | 0.0025 | 0.095 | 2 | 5 | 0.5 | 2.5 |
| Lim1215 |  | EGFR | 1250 cells | NA | 40000 cells | 0.04 | 0.06 | 0.0006 | 0.075 | 5 | 3 | 0.5 | 2.5 |
| Cell Line Name | Tissue of Origin | Driver | Plating density (384-well plates) | Plating density (96-well plates) | Plating density (6-well plates) | HER2i (Lapatinib) | PI3Ki (BKM120) | AKTi (MK2206) | mTORi (AZD8055) | HER2i (Lapatinib) | PI3Ki (BKM120) | AKTi (MK2206) | mTORi (AZD8055) |
| SKBR3 | Breast | HER2 | 2500 cells | NA | 100000 cells | 0.025 | 0.3 | 0.05 | 0.025 | 2.5 | 2.5 | 2.5 | 2.5 |
| HCC1954 |  | HER2 | 500 cells | NA | 40000 cells | 0.0125 | 0.4 | 0.5 | 0.025 | 5 | 3 | 5 | 2.5 |
| Organoid Name | Tissue of Origin | Driver | Plating density (384-well plates) | Plating density (96-well plates) | Plating density (6-well plates) | EGFRi (Gefitinib) | RAFi (LY3009120) | MEKi (Trametinib) | ERKi (SCH772984) | EGFRi (Gefitinib) | RAFi (LY3009120) | MEKi (Trametinib) | ERKi (SCH772984) |
| ITO034T | Lung | EGFR | NA | NA | NA | 0.065 | 0.035 | 0.0003 | 0.004 | NA | NA | NA | NA |
| TUM09 | CRC | EGFR | NA | NA | NA | 0.125 | 0.76 | 0.014 | 0.225 | NA | NA | NA | NA |
| TUM10 |  | KRAS | NA | NA | NA | 2.2 | 0.16 | 0.003 | 0.05 | NA | NA | NA | NA |

**Supplemental Table 2:** Compendium of patient, tumour, treatments and mutations information for all the PDXs used in the study.

|  | PDX name |  |  |  |
| --- | --- | --- | --- | --- |
|  | PDX-1 | PDX-2 | PDX-3 | PDX-4 |
| Age | 46 | 69 | 44 | 39 |
| Gender | Female | Male | Female | Female |
| Smoking Info | Former | Smoker | Former | Never |
| Diagnostic | Lung Adenocarcinoma | Lung Adenocarcinoma | Lung Adenocarcinoma | Lung Adenocarcinoma |
| EGFR mutation | Exon 19 deletion | Exon 19 deletion | L858R | Exon 19 deletion |
| Treatment 1 (T1) | Erlotinib | Erlotinib | Erlotinib | Afatinib |
| Alterations after T1 | T790M positive | MET amplification | MET overexpression | Unknown |
| Treatment 2 | Osimertinib | Gefitinib + Capmatinib | Gefitinib + Capmatinib | CBDCA + Pemetrexed |
| Treatment 3 | Osimertinib + Capmatinib | CDDP + Pemetrexed | Carboplatin + Gemcitabine + Nivolumab | CDDP + VP-16 |
| Source of PDX | After progressing to 2 lines | After progressing to 3 lines | After progressing to >3 lines | After progressing to 2 lines |
| Known alterations in the PDX | EGFR (T790M) and KRAS G12C | EGFR (del19) and METamp | EGFR (L858R) and METamp | TP53 mutation |

